# A non-invasive method to generate induced pluripotent stem cells from primate urine

**DOI:** 10.1101/2020.08.12.247619

**Authors:** Johanna Geuder, Mari Ohnuki, Lucas E. Wange, Aleksandar Janjic, Johannes W. Bagnoli, Stefan Müller, Artur Kaul, Wolfgang Enard

## Abstract

Comparing the molecular and cellular properties among primates is crucial to better understand human evolution and biology. However, it is difficult or ethically even impossible to collect matched tissues from many primates, especially during development. An alternative is to model different cell types and their development using induced pluripotent stem cells (iPSCs). These can be generated from many tissue sources, but non-invasive sampling would decisively broaden the spectrum of non-human primates that can be investigated. Here, we report the generation of primate iPSCs from urine samples. We first validate and optimize the procedure using human urine samples and show that Sendai virus transduction of reprogramming factors into urinary cells efficiently generates integration-free iPSCs, which maintain their pluripotency under feeder-free culture conditions. We demonstrate that this method is also applicable to gorilla and orangutan urinary cells isolated from a non-sterile zoo floor. We characterize the urinary cells, iPSCs and derived neural progenitor cells using karyotyping, immunohistochemistry, differentiation assays and RNA-sequencing. We show that the urine-derived human iPSCs are indistinguishable from well characterized PBMC-derived human iPSCs and that the gorilla and orangutan iPSCs are well comparable to the human iPSCs. In summary, this study introduces a novel and efficient approach to generate iPSCs non-invasively from primate urine. This will allow to extend the zoo of species available for a comparative approach to molecular and cellular phenotypes.

**Graphical Abstract:** Workflow overview for establishing iPSCs from primate urine
**(A)** We established the protocol for human urine based on a previous description (Zhou 2012). We tested volume, storage and culture conditions for primary cells and compared reprogramming by overexpression of OCT3/4, SOX2, KLF4 and MYC (OSKM) via lipofection of episomal vectors and via transduction of a sendai virus derived vector (SeV). **(B)** We used the the protocol established in humans and adapted it for unsterile floor-collected samples from non-human primates by adding Normocure to the first passages of primary cell culture and reprogrammed visually healthy and uncontaminated cultures using SeV. **(C)** Pluripotency of established cultures was verified by marker expression, differentiation capacity and cell type classification using RNA sequencing.

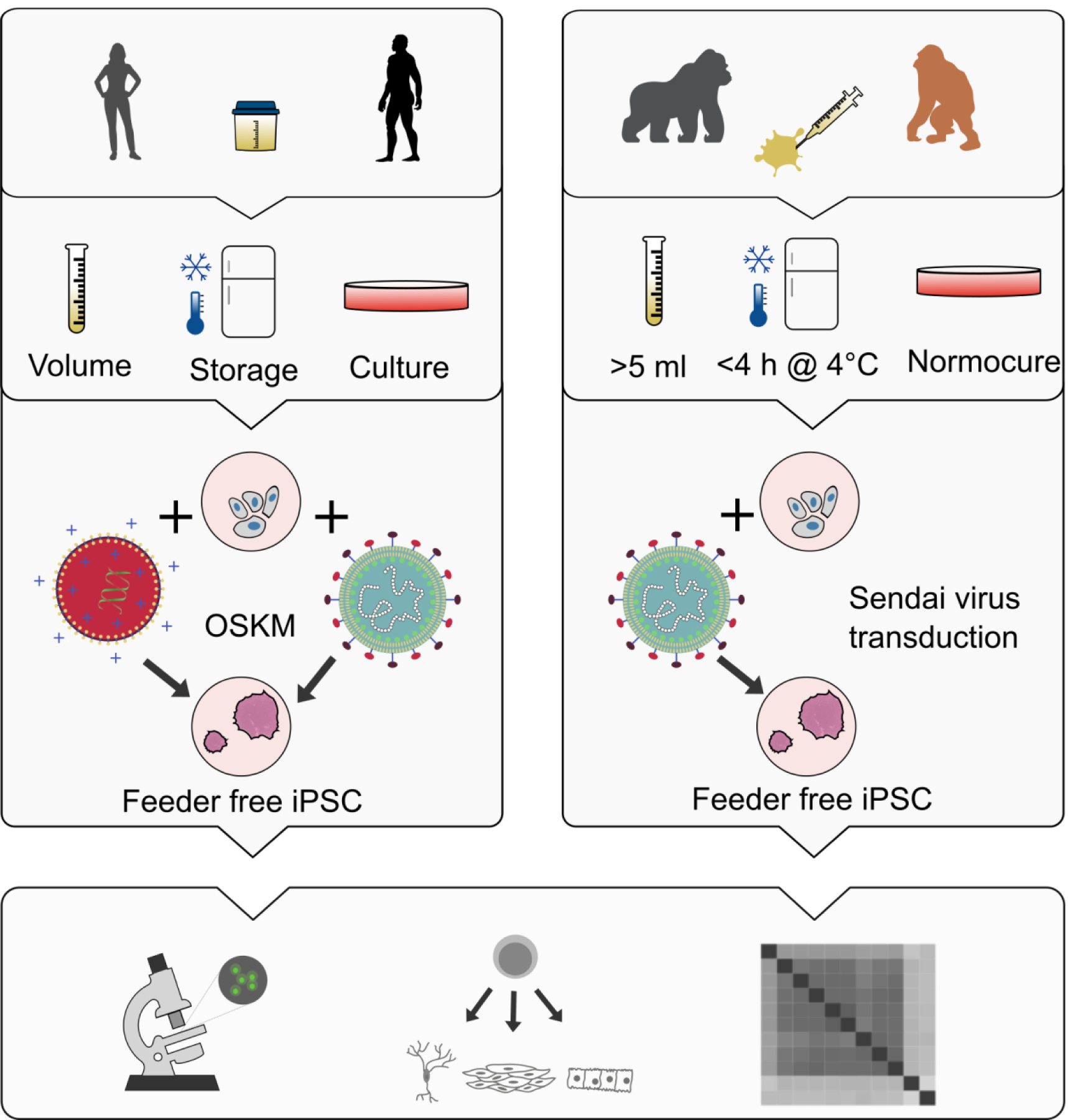

## Introduction

Primates are our closest relatives and hence play an essential role in biology, ecology and medicine. We share the vast majority of our genetic information and yet have considerable molecular and phenotypic differences (Pecon-Slattery, 2014). Understanding this genotype-phenotype evolution is crucial to understand the molecular basis of human-specific traits. Additionally, it is biomedically highly relevant to interpret findings made in model organisms, such as the mouse and to identify the conservation and functional relevance of molecular and cellular circuitries (Enard, 2012, 2016). However, obtaining comparable samples from different primates, especially during development, is practically and - more importantly - ethically very difficult or impossible.

Embryonic stem cells have the potential to partially overcome this limitation by their ability to differentiate into all cell types *in vitro* and divide indefinitely (Evans and Kaufman, 1981). However, the necessary primary material collection from an embryo is also practically and ethically difficult or impossible. Fortunately, a pluripotent state can also be induced in somatic cells by ectopically expressing four genes (Takahashi and Yamanaka, 2006). Since this discovery of induced pluripotency, great efforts have been made to identify suitable somatic cells (Raab et al., 2014) and optimize reprogramming methods (Schlaeger et al., 2015). Most of this research focused on human or mouse and while the methods are in general transferable and iPSCs from several different non-human primates (Gallego Romero et al., 2015; Nakai et al., 2018; Wunderlich et al., 2014) and other mammals (Ezashi et al., 2016; Stanton et al., 2018) have been generated, these methods have not been optimized for non-model organisms.

One major challenge for establishing iPSCs of various non-human primates is the acquisition of the primary cells. So far iPSCs have been generated from fibroblasts, peripheral blood cells or vein endothelial cells derived from postmortem tissue or during medical examinations (Gallego Romero et al., 2015; Nakai et al., 2018; Wunderlich et al., 2014); (Fujie et al., 2014; Morizane et al., 2017). However, also these sources impose practical and ethical constraints and therefore limit the availability of the primary material.

To overcome these limitations, we adapted a method of isolating reprogrammable cells from human urine samples (Zhou et al., 2011, 2012) and applied it to non-human primates (Figure 1). We find that primary cells can be isolated from unsterile urine sampled from the floor, can be efficiently reprogrammed using the integration-free Sendai Virus (Fusaki et al., 2009) and can be maintained under feeder-free conditions as shown by generating iPSCs from human, gorilla and orangutan.

**Figure 1.**
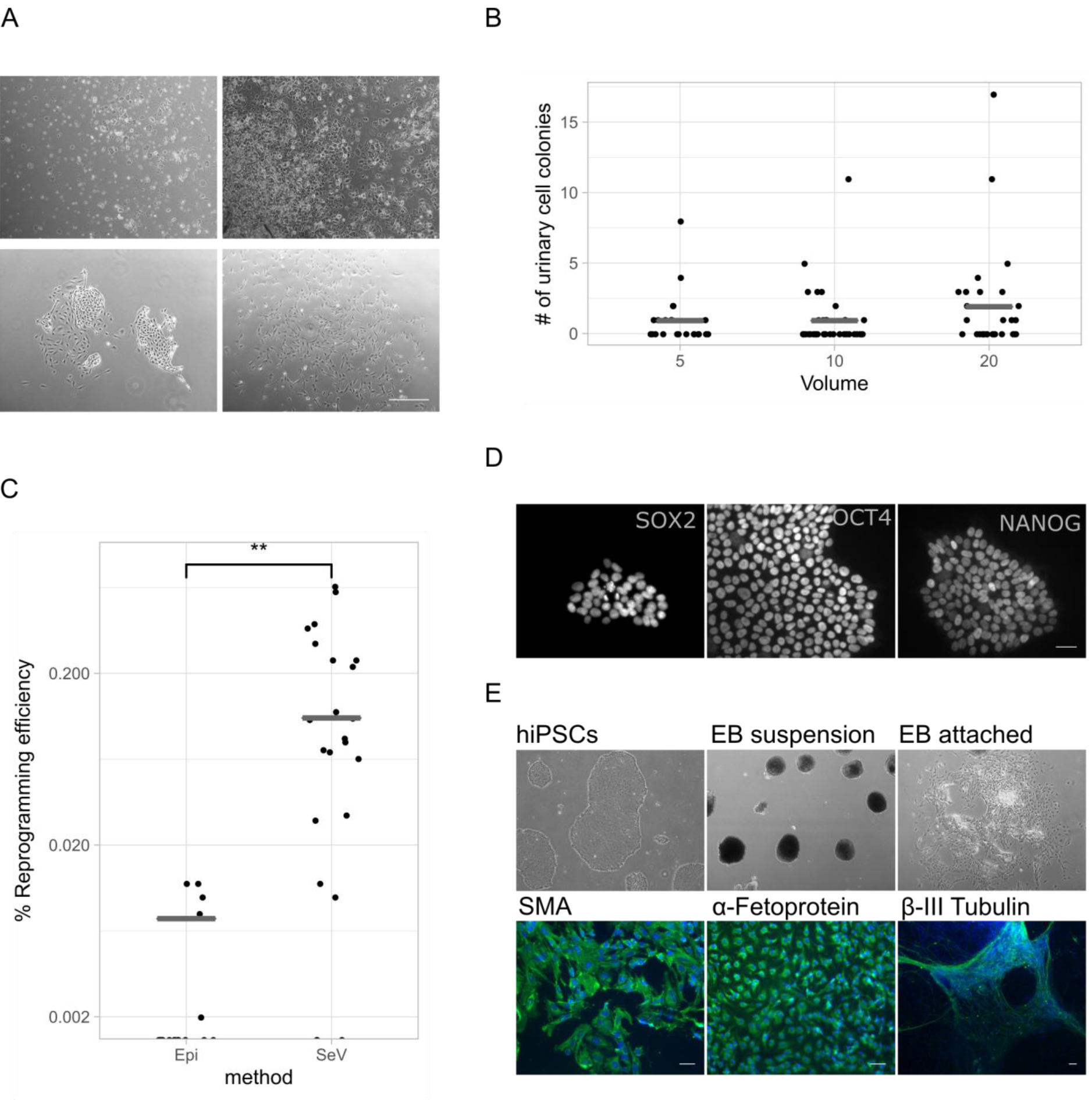
Establishing urinary cell isolation and reprogramming to iPSCs in human samples. **(A)** Human urine mainly consists of squamous cells and other differentiated cells that are not able to attach and proliferate (upper row). After ∼5 days, first colonies become visible and two types of colonies can be distinguished as described in Zhou (2012). Scale bars represent 400 μm **(B)** Isolation efficiency of urine varies between samples. The efficiency between 5 ml, 10 ml and 20 ml of starting material is not different (Fisher’s exact test *p* > 0.5) **(C)** SeV mediated reprogramming showed significantly higher efficiency than Episomal plasmids (Wilcoxon rank sum test: p=1.1e-05). **(D**,**E)** Established human colonies transduced with SeV expressed Nanog, Oct4 and Sox2; Scale bars represent 50 μm and **(F)** differentiated to cell types of the three germ layers; scale bar represents 100 µm. See also Figure S1.

## Results

### Isolating human urinary cells from small and stored samples

To assess which method is most suitable for isolating and reprogramming primate cells, we first tested different procedures using urinary cells from human samples. We collected urine from several humans in sterile beakers and processed them as described in Zhou et al (2011). As previously reported (Bharadwaj et al., 2013), we observed two morphologically distinct colony types that were indistinguishable after the first passage; the colonies consisted of grain-shaped cells that proliferated extensively (Figure 1A). In total we processed 19 samples of several individuals in 122 experiments using different volumes and storage times (Table S1). Per sample, we isolated an average of 7.6 colonies per 100 mL of urine when processing samples immediately. Isolation efficiencies varied considerably across samples (range: 0-70 colonies per 100 mL, Figure 1B), but did not differ between sexes. Furthermore, storing samples for up to 4 hours at room temperature or on ice did not influence the number of isolated colonies (9 samples, 7.4 colonies on average per 100 mL, range: 0-17). As sample volumes can be small for non-human primates, we also tested whether colonies can be isolated from 5, 10 or 20 mL of urine (Figure 1B). We found no evidence that smaller volumes have lower success rates as we found that for 42% of the 5 mL samples, we could isolate at least one colony (Table S1). Many more samples and conditions would be needed to better quantify the influence of different parameters on the isolation efficiency of colonies. However, of practical relevance is that small urine samples stored for a few hours at room temperature or on ice are a possible source to generate iPSCs from non-human primates.

### Reprogramming human urinary cells is efficient when using Sendai Virus transduction

Next, we investigated which integration-free overexpression strategy to induce pluripotency would be the most suitable for the isolated urine cells. To this end we compared transduction by a vector derived from the RNA-based Sendai Virus (Fujie et al., 2014; Fusaki et al., 2009) in suspension (Nakai et al., 2018), to lipofection with episomal plasmids (Epi) derived from the Eppstein Barr virus (Okita et al., 2011, 2013). Both systems have been previously reported to sufficiently induce reprogramming of somatic cells without the risk of genome integrations. Transduction of urinary cells with a Sendai Virus (SeV) vector containing Emerald GFP (EmGFP) showed substantially higher efficiencies than lipofection with an episomal plasmid (∼97% versus ∼20% EmGFP+; Figure S2A and S2B). We assessed the reprogramming efficiency of these two systems by counting colonies with a pluripotent-like cell morphology. Using SeV vectors, 0.19% of the cells gave rise to such colonies (Figure 1C). In contrast, using Episomal plasmids, only 0.009% of the cells gave rise to colonies with pluripotent cell-like morphology (N=23 and 18, respectively; Wilcoxon rank sum test: p=0.00005), resulting in at least one colony in 87% and 28% of the cases. Furthermore, the first colonies with a pluripotent morphology appeared 5 days after SeV transduction and 14 days after Epi lipofection. To test whether the morphologically defined pluripotent colonies also express molecular markers of pluripotency, we isolated flat, clear-edged colonies from 5 independently transduced urinary cell cultures on day 10. All clones expressed POU5F1 (OCT3/4), SOX2, NANOG and TRA-1-60 and differentiated into the three germ layers during embryoid body formation as shown by immunocytochemistry (Figure 1D, 1E). Notably, while the transduced cells also expressed the pluripotency marker SSEA4, this was also true for the primary urinary cells (Figure S2C). SSEA4 is known to be expressed in urine derived cells (Bharadwaj et al., 2013; Zhang et al., 2008) and hence it is an uninformative marker to assess the reprogramming of urinary cells to iPSCs. Clearance of SeV after transduction was routinely confirmed after the first five passages and the pluripotent state could be kept for over 100 passages (data not shown).

In summary, we find that the generation of iPSCs from human urine samples is possible from small volumes and most efficiently done using SeV transduction. Hence, we used this workflow for generating iPSCs from non-human primate cells.

### Isolating cells from unsterile primate urine

A decisive difference when sampling urine from non-human primates (NHPs) is the collection procedure. Samples from chimpanzees, gorillas and orangutans were collected by zoo keepers directly from the floor, often with visible contamination. Initially, culturing these samples was not successful due to the growth of contaminating bacteria. The isolation and culture of urinary cells only became possible upon the addition of Normocure (Invivogen), a broad-spectrum antibacterial agent that actively eliminates Gram+ and Gram-bacteria from cell cultures. We confirmed that Normocure did not affect the number of colonies isolated from sterile human samples (Table S1). Furthermore, many NHP samples also had volumes below 5 mL. While we tried to isolate cells from a total of 70 samples, only 24 samples had conditions verified for human urine as described above (>= 5 mL of sample, < 4h storage at RT or 4 °C and no visible contamination). From chimpanzees, gorillas and orangutans we collected a total of 87, 70 and 39 mL of urine in 11, 8 and 5 samples from several individuals and isolated 0, 5 and 2 colonies respectively (Table S2). For gorilla and orangutan this rate (7.3 and 5.2 colonies per 100 mL urine) is not significantly different from the rate found for human samples (6.0 per 100 mL across all conditions in Table S1, p=0.8 and 0.6, respectively, assuming a Poisson distribution). However obtaining zero colonies from 87mL of chimpanzee urine is less than expected given the rate found in human samples (p=0.005). So while isolating primary cells from urine samples seems comparable to human in two great ape species, it seems to have at least a 2-3 fold lower rate in our closest relatives, suggesting that the procedure might work in many, but not in all NHPs. Fortunately, it is possible to culture many samples in parallel so that screening for urinary cells in a larger volume with more samples is relatively easy.

The first proliferating cells from orangutan and gorilla could be observed after six to ten days (Figure 2A) in culture and could be propagated for several passages, as with the human cells. While we observed different proliferation rates and morphologies among samples, these did not differ systematically among individuals or species (Figure 2B). Infection with specific pathogens, including simian immunodeficiency virus (SIV), Herpes-B virus (BV), simian T cell leukemia virus (STLV) and simian betaretrovirus (SRV), was not detected in these cells (data not shown).

**Figure 2.**
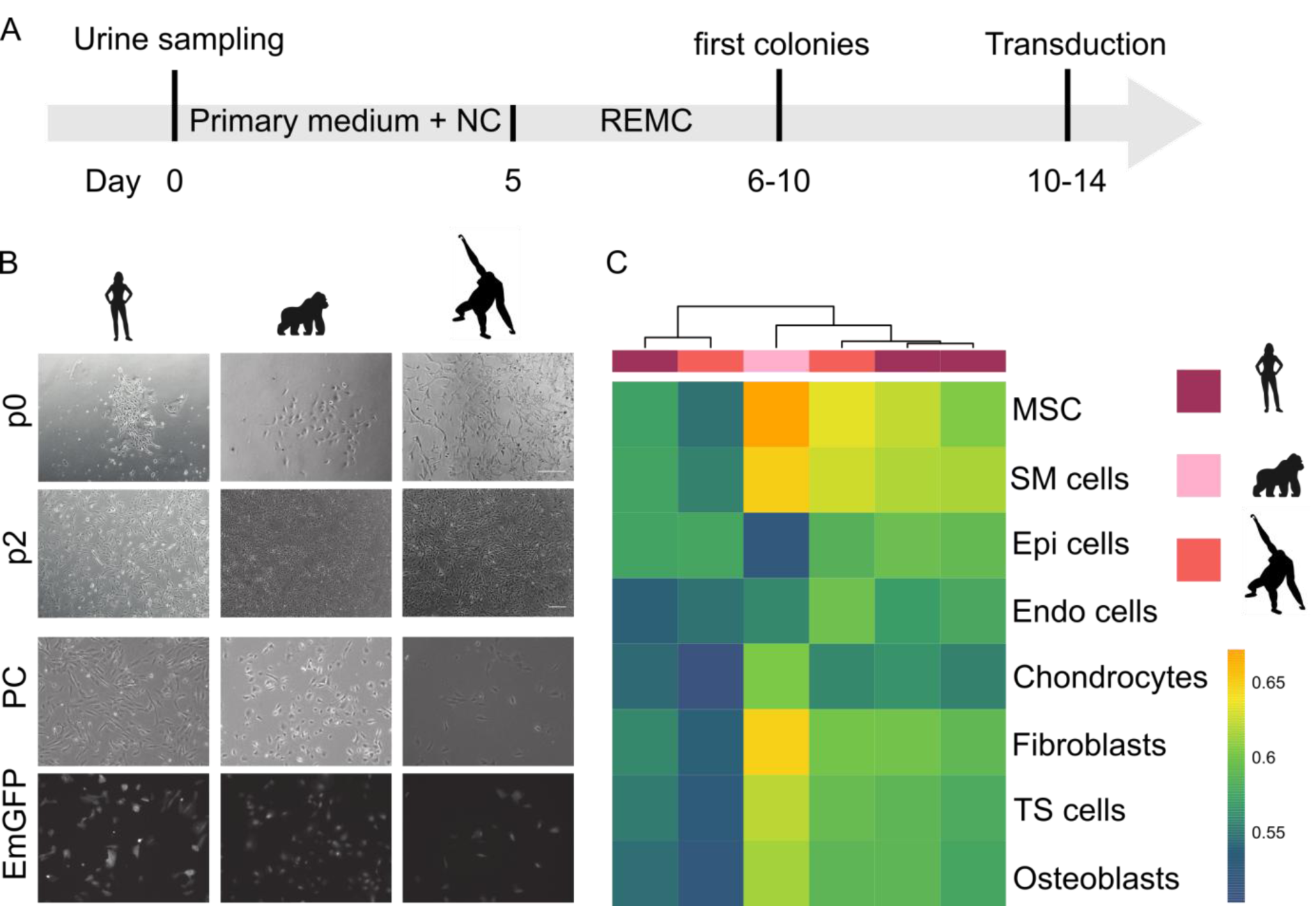
Isolation and characterization of primate urinary cells. **(A)** Workflow of cell isolation from primate urine samples. **(B)** Primary cells obtained from human, gorilla and orangutan samples are morphologically indistinguishable and display similar EmGFP transduction levels. Scale bars represent 400μm **(C)** The package SingleR was used to correlate the expression profiles from six samples of primate urinary cells (passage 1-3) to a reference set of 38 human cell types. The eight cell types with the highest correlations are shown (MSC = mesenchymal stem cells; SM = smooth muscle; Epi = epithelial; Endo = endothelial). Colour bar indicates correlation coefficients. See also Figure S2.

### Expression patterns of urinary cells are most similar to mesenchymal stem cells, epithelial cells and smooth muscle cells

To characterize the isolated urinary cells, we generated expression profiles using a 3’ tagged RNA-seq protocol (Bagnoli et al., 2018; Christoph Ziegenhain et al., 2017; Soumillon et al., 2014) on early passage primary urinary cells (p1-3) from three humans, one gorilla and one orangutan. Note that some of these samples contained cells from 1-4 different colonies (Table S1 and S2) and hence could be mixtures of different cell types. To classify these urinary cells we compared their expression profile to 713 microarray expression profiles grouped into 38 cell types (Mabbott et al., 2013) using the SingleR package (Aran et al., 2019). SingleR uses the most informative genes from the reference dataset and iteratively correlates it with the expression profile to be classified. The most similar cell types were mesenchymal stem cells, epithelial cells and/or smooth muscle cells and at least two groups are evident among the six samples (Figure 2C). One group of four samples shows relatively high similarity to mesenchymal cells and smooth muscle cells, in agreement with a characterization of urine-derived stem cells as being similar to mesenchymal stem cells with respect to surface markers and differentiation capacity (Bharadwaj et al., 2013). The other group with one human and one orangutan sample (derived from two and one colonies, respectively) is equally similar to mesenchymal stem cells, epithelial cells and smooth muscle cells. These relatively similar correlation coefficients might indicate that the exact cell types present in our samples might not be among the 38 cell types in the reference data. Better reference data like those generated within the Human Cell Atlas (Rozenblatt-Rosen et al., 2017) will be needed to characterize the diversity of urine-derived cells more thoroughly and more systematically. In the context of this study, it is relevant that our findings are in general agreement with the notion that several different types of proliferating cell types can be isolated from shed human urine (Bento et al., 2020), and that the same types of cells are found - as expected - in primates.

### Reprogramming efficiency of urinary cells is similar in humans and other primates

To generate iPSCs from the urinary cells isolated from gorilla and orangutan, we used Sendai Virus (SeV) transduction that we found to be efficient for human urinary cells (Figure 3A). Human, gorilla and orangutan urinary cells showed similarly high transduction efficiencies with the EmGFP SeV vector (data not shown). Transduction with the reprogramming SeV vectors led to first morphological changes already after 2 days in all three species, when cells began to form colonies and became clearly distinguishable from the primary cells (Figure 3B). When flat, clear-edged colonies appeared that contained cells with a large nucleus to cytoplasm ratio, these colonies were picked and plated onto a new dish. We found that the efficiency and speed of reprogramming was variable (Figure S2B), probably depending on the cell type, the passage number and the acute state (“health”) of the cells, in concordance with the variability and efficiency found in other studies utilizing urine cells as source for iPSCs (Zhou et al., 2011). Also the mean reprogramming efficiency over all replicates was different (Kruskal-Wallis Test, p= 0.015) for human (0.19%) and gorilla (0.28%) and orangutan (0.061%). However, many more samples would be necessary to disentangle the effects of all these factors. Of note, we observed that the orangutan iPSCs showed more variability in proliferation rates and morphology compared to human and gorilla iPSCs. Several subcloning steps were needed until a morphologically stable clone could be generated. However, the resulting iPSCs were stable and had the same properties as the other iPSCs (see below). To what extent this is indeed a property of the species is currently unclear. Importantly, from all primary samples that were transduced, colonies with an iPSC morphology could be obtained. So while considerable variance in reprogramming exists, the reprogramming efficiency is sufficiently high and sufficiently similar in humans, gorillas and orangutans.

**Figure 3.**
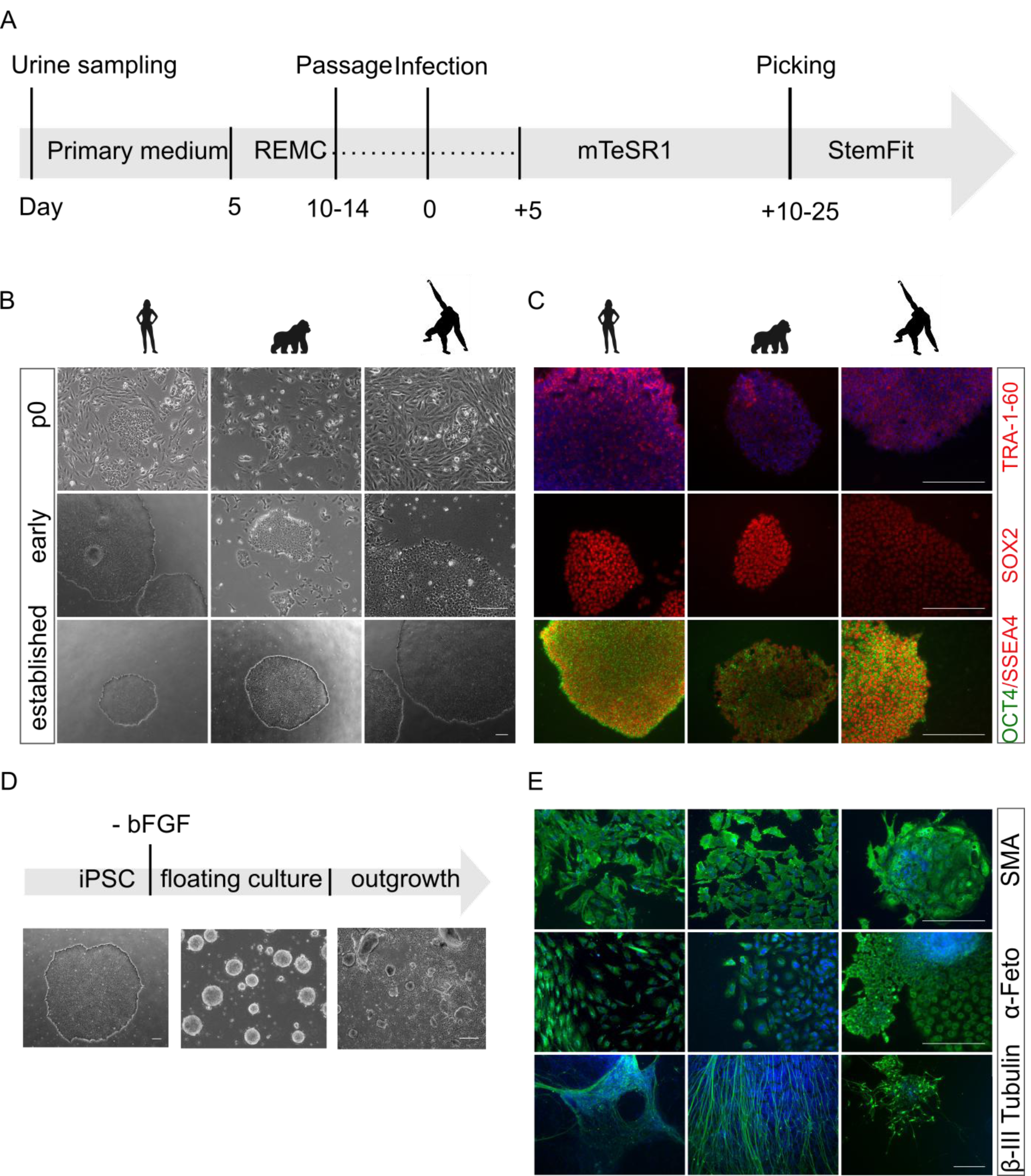
Generation and characterization of primate iPSCs. **(A)** Workflow for reprogramming of primate urinary cells. **(B)** Cell morphology of the three species is comparable before (p0), during (p1-3) and after reprogramming (∼p5). **(C)** Immunofluorescence analysis of pluripotency associated proteins at passage X-Y: TRA-1-60, SSEA4, OCT4 and SOX2. Nuclei were counterstained with DAPI. **(D)** Differentiation potency into the three germ layers. iPSC colony before differentiation, after 8 days of floating culture and after 8 days of attached culture. **(E)** Immunofluorescence analyses of ectoderm (β-III Tubulin), mesoderm (α-SMA) and endoderm markers (α-Feto) after EB outgrowth. Nuclei were counterstained with DAPI. Scale bars represent 400μm. See also Figure S3 & Figure S4.

### Urine derived primate iPSCs are comparable to human iPSCs

We could generate at least two lines per individual from all primary cell samples that all showed Oct3/4, TRA-1-60, SSEA4, SOX2 and NANOG immunofluorescence (Figure 3C). Furthermore, 5 out of 5 tested iPSC lines showed a normal karyotype, without chromosomal alterations (Figure S4). iPSCs from all species could be expanded for more than fifty passages, while maintaining their pluripotency, as shown by pluripotency marker expression (Figure 3C) and differentiation capacity via embryoid body formation (Figure 3E). Both the human and NHP iPSCs differentiated into ectoderm (beta-III Tubulin), mesoderm (α-SMA) and endoderm (AFP) lineages (Figure 3E). Dual-SMAD inhibition led to the formation of neurospheres in floating culture, as confirmed by neural stem cell marker expression (NESTIN+, PAX6+) using qRT-PCR (Figure S3).

To further assess and compare the urine-derived iPSCs, we generated RNA-seq profiles from nine human, three gorilla and four orangutan iPSC lines as well as the six corresponding primary urinary cells (see analysis above). As an external reference, we added a previously reported and well characterized blood-derived human iPS cell line that was generated using episomal vectors and adapted to the same feeder-free culture conditions as our cells (1383D2) (Nakagawa et al., 2014). All lines were grown and processed under the same conditions and in a randomized order in one experimental batch. We picked one colony per sample and used a 3’ tagged RNA-seq protocol (Bagnoli et al., 2018; Christoph Ziegenhain et al., 2017; Soumillon et al., 2014) to generate expression profiles with 19,000 genes detected on average.

We classified the expression pattern of the iPSCs relative to the reference dataset of 38 cell types using SingleR as described for the urinary cells. ES cells or iPS cells are clearly the most similar cell type for all our iPS samples including the external PBMC-derived iPSC line (Figure 4A). Principal component analysis of the 500 most variable genes (Figure 4B), shows clear clustering of the samples according to cell type (54% of the variation in PC1) and species (23% of the variance in PC2). The external, human blood-derived iPSC line is interspersed among our human urine derived iPS cell lines. Using the pairwise Euclidean distances between samples to assess similarity, they also cluster first by cell type and then by species (Figure S2D). When classifying the expression pattern of the iPSCs relative to a single cell RNA-seq dataset covering distinct human embryonic stem cell derived progenitor states (Chu et. al. 2016), again all our iPSC lines are most similar to embryonic stem cells and are indistinguishable from the external PBMC-derived iPSC line (Figure 4C). Finally, expression distances within iPS cells of the same species were similar, independent of the individual and donor cell type (Figure 4D).

**Figure 4.**
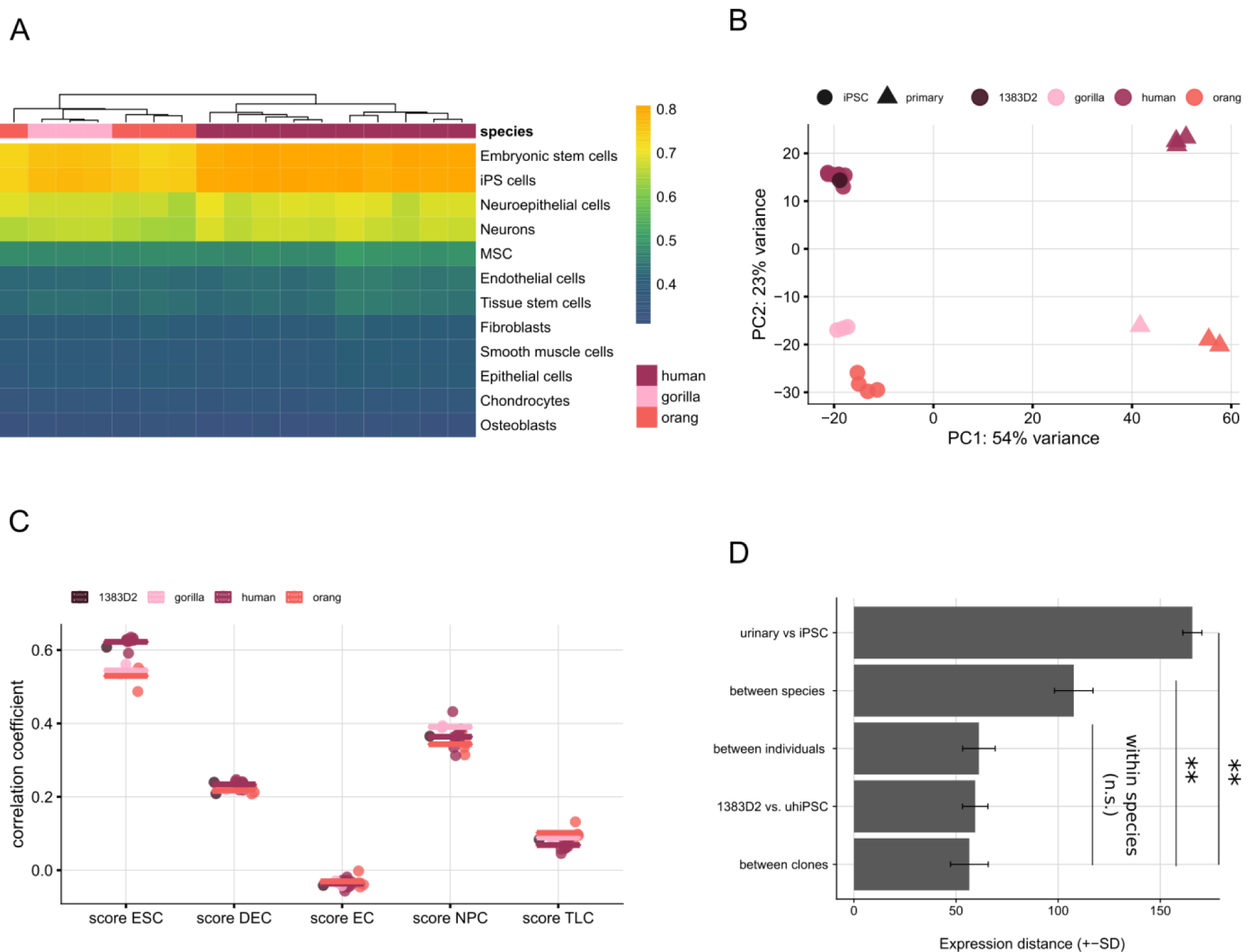
Characterization of primate iPSCs by expression profiling. **(A)** The package SingleR was used to correlate the expression profiles from seventeen samples of primate iPSCs (passage 1-3) to a reference set of 38 human cell types. The twelve cell types with the highest correlations are shown (MSC = mesenchymal stem cells). All lines are similarly correlated to embryonic stem cells and iPS cells. Colour bar indicates correlation coefficients. **(B)** Principal component analysis of primary cells and derived iPSC lines using the 500 most variable genes. PC1 separates the cell types and PC2 separates the species from each other **(C)** Correlation coefficient of iPSCs compared to a single cell dataset covering distinct human embryonic stem cell derived progenitor states (Chu et. al. 2016). **(D)** Expression distances of all detected genes are averaged from pairwise distances for six different groups of comparisons. Note that the distance between and among individuals and species is calculated within iPSCs and distances between individuals within species. Pairwise t-tests are all below 0.01 (**) for comparisons to the cell-type and species distance and all above 0.05 (n.s.) for comparisons within the species. See also Figure S2.

Taken together, these analyses do not only indicate that our urine derived iPS cells show a pluripotent expression profile and differentiate as expected for iPS cells but can also not be distinguished from an iPSC line derived in another laboratory from another cell type with another vector system. Hence, the expression differences among species are far larger than these technical sources of variation, indicating that these cells are well suited to assess species differences among primates.

## Discussion

Here, we adapted a previously described protocol (Zhou et al., 2012) to isolate proliferating cells from unsterile primate urine. We show that these urinary cells can be efficiently reprogrammed into integration-free and feeder-free iPSCs, which areclosely comparable among each other and to other iPSCs.

Our findings have implications for generating and validating iPSCs from primates and other species for comparative studies. Additionally, some aspects might also be of relevance when generating iPSCs from human urinary cells for medical studies.

### Implications for using urine for generating human iPSCs

Human urine mainly contains cells, such as squamous cells, which are terminally differentiated and cannot attach or proliferate in culture. The first proliferating cells from human urine were isolated in 1972 (Sutherland and Bain, 1972) and since then a variety of different cells have been isolated and described that can proliferate, differentiate and be reprogrammed to iPSCs (see (Bento et al., 2020) for a recent overview). As these urine-derived stem cells (UDSCs) can be isolated non-invasively at low costs and reprogram efficiently (Zhou et al., 2012), they are increasingly used to generate iPSCs from patients (e.g. (Ernst, 2020; Gaignerie et al., 2018; Xue et al., 2013). Perhaps the only major drawback of using UDSCs for iPSC generation is that the number of UDSCs per millilitre is quite variable within and among individualsEven when collecting large volumes, isolation can fail as we also see in our study (Table S1).

The quantitative relation between morphology, marker expression, potency and reprogramming efficiency among the different UDSCs is not clear. The six RNA-seq profiles of UDSC colonies from humans and apes presented here are to our knowledge the first genome-wide expression patterns from UDSCs. We find that four colonies have an expression profile correlating best with mesenchymal stem cells and two colonies have a profile that matches equally well to epithelial, smooth muscle and mesenchymal stem cells (Fig. 2C). So we find at least two different cell types, of which at least one is not among a comprehensive reference cell type data set. This is generally in agreement with previous studies (Bharadwaj et al., 2013; Zhang et al., 2008), but more comprehensive studies including comparisons to reference cell types that will be generated within the Human Cell Atlas (Rozenblatt-Rosen et al., 2017) will be needed to better characterize the diversity of UDSCs. For the purpose of using these cells for iPSC generation, these differences seem to not matter much, as all types seem to reprogram with sufficient efficiency. Regarding the reprogramming method, we find that transduction using the commercial Sendai Virus based vector in suspension (Nakai et al., 2018) is substantially more efficient for UDSCs than lipofection of episomal plasmids and also leads to a very quick change of morphology already within 2 days. While it is established that Sendai Virus reprogramming is an expensive but efficient method to generate iPSCs from fibroblasts (Churko et al., 2017; Schlaeger et al., 2015), our findings indicate that especially the suspension method might be the most efficient method for UDSCs. Finally, a relevant side note of our findings is that SSEA4, which is sometimes used as a marker for pluripotency (Pera et al., 2000; Thomson et al., 1998), is not useful when starting from urinary cells as these express SSEA4 already at high levels (Figure S1C). In summary, our findings inform a few aspects of generating iPSCs from human urine. However, their main relevance is for generating iPSCs from other species.

### Implications for isolating urinary cells from mammals

Our results have implications for isolating urinary stem cells for the generation of iPSC from primates and other mammals. This could be useful in contexts where invasive sampling is difficult, as it is the case for non-model primates and mammals, and where iPSCs are needed for conservation (Stanton et al., 2018) or comparative approaches (see below). So how likely is it that one can find UDSCs also in other primates and mammals? In humans, many UDSCs originate from the kidney as shown in a female patient transplanted with a male kidney (Bharadwaj et al., 2013). We isolated UDSCs from orangutan and gorilla and found similar transcriptional profiles, morphologies and growth characteristics. However, our failure to isolate UDSCs from chimpanzees suggests that some species might have at least 2-3 times less of those cells in their urine. Given the general similarity of the urinary tract in mammals and our successful isolation of UDSCs in two apes, it seems likely that most primates, and maybe also most mammals, shed UDSCs in their urine. However, it is also likely that the concentration of these cells varies even among closely related species as our results from chimpanzees suggest. Hence, from which species UDSCs can be isolated in practice will depend on this concentration and the available amount of urine. Fortunately, the cost-efficiency of the culturing system allows that this can be easily tested for a given species. Furthermore, our procedure to use unsterile samples from the floor to isolate such cells, broadens the practical implementation of this approach considerably.

### Implications for generating iPSCs in many species

Given that it is possible to isolate UDSCs from a species, the efficiency of reprogramming and iPSC maintenance will determine whether one can generate stable iPSCs from them. Fortunately, the efficiency of reprogramming UDSCs is high, probably higher than for many other primary cell types (Raab et al., 2014). This is especially true when using SeV transduction in suspension as is evident from the fact that we could generate iPSCs from all twelve UDSC reprogramming experiments (Table S3). To what extent this reprogramming procedure works in other species is unclear. The Sendai virus is promiscuous and infects all mammalian cells (Nishimura et al., 2011). Pluripotent stem cells are also present in blastocysts of all mammals and iPSCs have been generated from many species (Stanton et al., 2018). Using human reprogramming factors and culture conditions has led to the generation of iPSCs from many mammals and even avian species (Stanton et al., 2018), albeit with over 10-fold lower reprogramming efficiencies (Ben-Nun et al., 2011; Stauske et al., 2020). So while in principle it should be possible to isolate iPSCs from many or even all mammals, variation in reprogramming efficiency with human factors and culture conditions to keep cells pluripotent with and without feeder cells (Stauske et al., 2020) will considerably vary among species and will make it practically difficult to obtain and maintain iPSCs from some species. Investigating the cause of this variation more systematically will be important to better understand pluripotent stem cells in general and to generate iPSCs from many species in practice. Recent examples of such fruitful investigations include the optimization of culture conditions for baboons (Navara et al., 2018), and the optimization of feeder-free culture conditions for rhesus macaques and baboons (Stauske et al., 2020). A related aspect of generating iPSCs from different species is testing whether iPSCs from a given species are actually *bona fide* iPSCs. While for humans a variety of tools exist, such as predictive gene expression assays, validated antibody stainings and SNP arrays for chromosomal integrity, these tools can not be directly transferred to other species. Fortunately, due to the availability of genome sequences, RNA-sequencing in combination with human or mouse reference cell types to which generated iPSCs can be compared, the characterization of non-human iPSCs becomes feasible as also shown in this paper. In summary, while extending the zoo of comparable iPSCs is a daunting task and requires considerable more method development, we think our method to isolate UDSCs from unsterile urine could be a promising tool in this endeavor.

### Implications for a comparative approach to molecular phenotypes

So assuming that our approach works in at least some non-human primates (NHPs), the effectiveness and non-invasiveness of the protocol allows sampling many more individuals and species than currently possible. Why is this important? So far, iPSCs have been generated from a few individuals in a few NHP species. One main application is to model biomedical applications of iPSCs in primates such as rhesus macaques or marmosets (Hong et al., 2014). As these species are anyway used as model organisms, non-invasive sampling is less of an issue. Another main application are studies of the type “what makes us humans?” by comparing molecular phenotypes like gene expression levels in humans, chimpanzees and an outgroup (Gallego Romero et al., 2015; Kanton et al., 2019; Marchetto et al., 2013; Wunderlich et al., 2014). As recently reminded (Kelley and Gilad, 2020), proper study design and comparability is crucial also for these studies and larger sample sizes are certainly needed to infer human-specific changes more robustly from iPSCs and their derivatives like brain organoids. A third type of question with considerable potential has been explored much less, namely using iPSCs in a comparative framework to identify molecular or cellular properties that are conserved, i.e. functional across species (Enard, 2012, 2016; Housman and Gilad, 2020).This is similar to the comparative approach on the genotype level in which DNA or protein sequences are compared in orthologous regions among several species to identify conserved, i.e. functional elements (Alföldi and Lindblad-Toh, 2013). This information is crucial for example when inferring the pathogenicity of genetic variants (Kircher et al., 2014). Accordingly, it would be useful to know whether a particular phenotypic variant, e.g. a disease associated gene expression pattern, is conserved across species. This requires to compare the orthologous cell types and states among several species. Primates are well suited for such an approach, because they bridge the evolutionary gap between human and its most important model organism, the mouse and because phenotypes and orthologous cell states can be more reliably compared in closely related species. However, for practical and ethical reasons, orthologous cell states are difficult to obtain from several different primates. Hence, just as human iPSCs allow to study cell types and states that are for practical and ethical reasons not accessible, primate iPSCs allow to extend the comparative approach to these cell types and states, leveraging unique evolutionary information that is not only interesting per se, but could also be of biomedical relevance. As our method considerably extends the possibilities to derive iPSCs from primates, it could contribute towards leveraging the unique information generated during millions of years of primate evolution.

## Experimental Procedures

### Data and Code availability

RNA-seq data generated here are available at GEO under accession number GSE155889. Code is available upon request.

### Experimental Model and Subject Details

#### Human urine samples

Human urine samples from 8 healthy volunteers were obtained with written informed consent and processed anonymously in accordance with the ethical standards of the responsible committee on human experimentation (20-122, Ethikkommission LMU München). Additional information on the samples are available in Table S1.

#### Primate urine samples

Primate urine was collected at the Hellabrunn Zoo in Munich, Germany. Caretakers took up available urin on the floor with a syringe, hence the collection procedure was fully non-invasive without additional perturbation of animals. From which individual the sample came was not evident during the collection due to the procedure. Additional information on the samples can be found in Table S2.

#### iPSC lines

iPSC lines were generated from human and primate urinary cells. Reprogramming was done using two different techniques. Reprogramming using SeV (Thermo Fisher) was performed as suspension transduction as described before (Nakai et al., 2018). Episomal vectors were transfected using Lipofectamine 3000 (Thermo Fisher). iPSCs were cultured under feeder-free conditions on Geltrex (Thermo Fisher) -coated dishes in StemFit medium (Ajinomoto) supplemented with 100 ng/mL recombinant human basic FGF (Peprotech), 100 U/mL Penicillin and 100 μg/mL Streptomycin (Thermo Fisher) at 37°C with 5% carbon dioxide. Cells were routinely subcultured using 0.5 mM EDTA. Whenever cells were dissociated into single cells using 0.5 × TrypLE Select (Thermo Fisher) or Accumax (Sigma Aldrich), the culture medium was supplemented with 10 µM Rho-associated kinase (ROCK) inhibitor Y27632 (BIOZOL) to prevent apoptosis.

### Method Details

#### Isolation of cells from urine samples

Urine from human volunteers was collected anonymously in sterile tubes. Usually a volume of 5 to 50 mL was obtained. Urine from NHPs was collected from the floor at Hellabrunn Zoo (Munich) by the zoo personnel, using a syringe without taking special precautions while collecting the samples. Samples were stored at 4°C until processing with a maximum time span of 5 hours. Isolation of primary cells was performed as previously described by Zhou et al. 2012. Briefly, the sample was centrifuged at 400 x g for 10 minutes and washed with DPBS containing 100 U/mL Penicillin, 100 μg/mL Streptomycin (Thermo Fisher), 2.5 µg/mL Amphotericin (Sigma-Aldrich). Afterwards, the cells were resuspended in urinary primary medium consisting of 10% FBS (Life Technologies), 100 U/mL Penicillin, 100 μg/mL Streptomycin (Thermo Fisher), REGM supplement (ATCC) in DMEM/F12 (TH. Geyer) and seeded onto one gelatine coated well of a 12-well-plate. To avoid contamination stemming from the unsanitary sample collection, 100 µg/mL Normocure (Invivogen) was added to the cultures until the first passage. 1 milliliter of medium was added every day until day 5, where 4 mL of the medium was aspirated and 1 mL of renal epithelial and mesenchymal cell proliferation medium RE/MC proliferation medium was added. RE/MC consists of a 50/50 mixture of Renal Epithelial Cell Basal Medium (ATCC) plus the Renal Epithelial Cell Growth Kit (ATCC) and mesenchymal cell medium consisting of DMEM high glucose with 10% FBS (Life Technologies), 2mM GlutaMAX-I (Thermo Fisher), 1x NEAA (Thermo Fisher), 100 U/mL Penicillin, 100 μg/mL Streptomycin (Thermo Fisher), 5 ng/mL bFGF (PeproTech), 5 ng/mL PDGF-AB (PeproTech) and 5 ng/mL EGF (Miltenyi Biotec). Half of the medium was changed every day until the first colonies appeared. Subsequent medium changes were performed every second day. Passaging was conducted using 0.5x TrypLE Select (Thermo Fisher). Typically 15×10^3^ to 30×10^3^ cells were seeded per well of a 12-well plate.

#### Generation of NHP iPSCs by Sendai virus vector infection

Infection of primary cells was performed with the CytoTune™-iPS 2.0 Sendai Reprogramming Kit (Thermo Fisher) at a MOI of 5 using a modified protocol. Briefly, 7×10^5^ urine derived cells were incubated in 100 µl of the CytoTune 2.0 SeV mixture containing three vector preparations: polycistronic Klf4–Oct3/4–Sox2, cMyc, and Klf4 for one hour at 37°C. To control transduction efficiency 3.5×10^5^ cells were infected with CytoTune-EmGFP SeV. Infected cells were seeded on Geltrex (Thermo Fisher) coated 12-well-plates, routinely 10×10^3^ and 25×10^3^ cells per well. Medium was replaced with fresh Renal epithelial and mesenchymal cell proliferation medium RE/MC (ATCC) every second day. On day 5, medium was changed to mTeSR1 (Stemcell Technologies), with subsequent medium changes every second day. After single colony picking, cells were cultured in StemFit (Ajinomoto) supplemented with 100 ng/mL recombinant human basic FGF (Peprotech), 100 U/mL Penicillin and 100 μg/mL Streptomycin (Thermo Fisher).

#### Immunostaining

Cells were fixed with 4% PFA, permeabilized with 0.3% Triton X-100, blocked with 5% FBS and incubated with the primary antibody diluted in 1% BSA and 0.3% Triton X-100 in PBS overnight at 4°C. The following antibodies were used: Human alpha-Smooth Muscle Actin (R&D Systems, MAB1420), Human/Mouse alpha-Fetoprotein/AFP (R&D Systems, MAB1368), Nanog (R&D Systems, D73G4), Neuron-specific beta -III Tubulin (R&D Systems, MAB1195), Oct-4 (NEB, D7O5Z), Sox2 (NEB, 4900S), SSEA4 (NEB, 4755), EpCAM (Fisher Scientific, 22 HCLC) and TRA-1-60 (Thermo Fisher, REA157). The next day, cells were washed and incubated with the secondary antibodies for one hour at room temperature. Alexa 488 rabbit (Thermo Fisher, A-11034) and Alexa 488 mouse (Thermo Fisher, A-21042) were used in a 1/500 dilution. Nuclei were counterstained using DAPI (Sigma Aldrich) at a concentration of 1 µg/mL.

#### Karyotyping

iPSCs at ∼80% confluency were treated with 50 ng/mL colcemid (Thermo Fisher) for 2 hours, harvested using TrypLE Select (Thermo Fisher) and treated with 75mM KCL for 20 minutes at 37°C. Subsequently, cells were fixed with methanol / acetic acid glacial (3:1) at - 20°C for 30 minutes. After two more washes of the fixed cell suspension in methanol/acetic (3:1), chromosome spreads were prepared using one drop of the fixed cell suspension.

#### RT-PCR and PCR analyses

Total RNA was extracted from cells lysed with Trizol using the Direct-zol RNA Miniprep Plus Kit (Zymo Research, R2072). 1 µg of total RNA was reverse transcribed using Maxima H Minus Reverse Transcriptase (Thermo Fisher) and 5 µM random hexamer primers. Conditions were as follows: 10 mins at 25°C, 30 mins at 50°C and then 5 mins at 85°C. Quantitative polymerase chain reaction (qPCR) studies were conducted on 5 ng of reverse transcribed total RNA in duplicates using PowerUp SYBR Green master mix (Thermo Fisher) using primers specific for NANOG, OCT4, PAX6 and NESTIN. Each qPCR consisted of 2 mins at 50°C, 2 mins at 95°C followed by 40 cycles of 15 s at 95°C, 15 s at 55°C and 1 min at 72°C. Cycle threshold was calculated by using default settings for the real-time sequence detection software (Thermo Fisher). For relative expression analysis the quantity of each sample was first determined using a standard curve and normalized to GAPDH and the average target gene expression (deltaCt/average target gene expression).

Genomic DNA for genotyping was extracted using DNeasy Blood & Tissue Kit (Qiagen). PCR analyses were performed using DreamTaq (Thermo Fisher). Primate primary cells were genotyped using primers that bind species-specific Alu insertions (adapted from (Herke et al., 2007).

To confirm the transgene-free status of the iPSC lines, SeV specific primers were used described in CytoTune®-iPS 2.0 Sendai Reprogramming Kit protocol (Thermo Fisher).

#### In vitro differentiation

For embryoid body formation iPSCs from one confluent 6-well were collected and subsequently cultured on a sterile bacterial dish in StemFit without bFGF. During the 8 days of suspension culture, medium was changed every second day. Subsequently, cells were seeded into six gelatin coated wells of a 6-well-plate. After 8 days of attached culture, immunocytochemistry was performed using α-fetoprotein (R&D Systems, MAB1368) as endoderm, α-smooth muscle actin (R&D Systems, MAB1420) as mesoderm and β-III tubulin (R&D Systems, MAB1195) as ectoderm marker.

For directed differentiation to neural stem cells (NSCs) cells were dissociated and 9×10^3^ cells were plated into each well of a low attachment U-bottom 96-well-plate in 8GMK medium consisting of GMEM (Thermo Fisher), 8% KSR (Thermo Fisher), 5.5 mL 100x NEAA (Thermo Fisher), 100mM Sodium Pyruvate (Thermo Fisher), 50mM 2-Mercaptoethanol (Thermo Fisher) supplemented with 500nM A-83-01 (Sigma Aldrich), 100nM LDN 193189 (Sigma Aldrich) and 30µM Y27632 (biozol). Half medium change was performed at d 4, 8, 11. Neurospheres were lysed in TRI reagent (Sigma Aldrich) at day 7 and differentiation was verified using qRT PCR.

#### Bulk RNA-seq library preparation

One colony per clone corresponding to ∼2×10^4^ cells and 2×10^3^ primary cells of each individual were lysed in RLT Plus (Qiagen) and stored at −80°C until processing. A modified SCRB-seq protocol was used for library preparation (Bagnoli et al., 2018; Christoph Ziegenhain et al., 2017; Soumillon et al., 2014). Briefly, proteins in the lysate were digested by Proteinase K (Ambion), RNA was cleaned up using SPRI beads (GE, 22%PEG). In order to remove isolated DNA, samples were treated with DNase I for 15 mins at RT. cDNA was generated by oligo-dT primers containing well specific (sample specific) barcodes and unique molecular identifiers (UMIs). Unincorporated barcode primers were digested using Exonuclease I (New England Biolabs). cDNA was pre-amplified using KAPA HiFi HotStart polymerase (Roche) and pooled before Nextera libraries were constructed from 0.8 ng of preamplified cleaned up cDNA. 3′ ends were enriched with a custom P5 primer (P5NEXTPT5, IDT) and libraries were size selected using a 2% E-Gel Agarose EX Gels (Life Technologies), cut out in the range of 300–800 bp, and extracted using the MinElute Kit (Qiagen) according to manufacturer’s recommendations.

#### Sequencing

Libraries were paired-end sequenced on an Illumina HiSeq 1500 instrument. Sixteen bases were sequenced with the first read to obtain cellular and molecular barcodes and 50 bases were sequenced in the second read into the cDNA fragment.

#### Data processing and analysis

All raw fastq data were processed with zUMIs (Parekh et al., 2018) using STAR 2.6.0a (Dobin et al., 2013) to generate expression profiles for barcoded UMI data. All samples were mapped to the human genome (hg38). Gene annotations were obtained from Ensembl (GRCh38.84). Samples were filtered based on number of genes and UMIs detected, and genes were filtered using HTS Filter. DESeq2 (Love et al., 2014) was used for normalization and variance stabilized transformed data was used for principal component analysis and hierarchical clustering.

Mitochondrial and ribosomal reads were excluded and singleR was used to classify the cells. SingleR was developed for unbiased cell type recognition of single cell RNA-seq data, however, here we applied the method to our bulk RNA seq dataset (Aran et al., 2019). The 200 most variable genes were used in the ‘de’ option of SingleR to compare the obtained expression profiles to (Chu et al., 2016) as well as HPCA (Mabbott et al., 2013). Based on the highest pairwise correlation between query and reference, cell types of the samples were assigned based on the most similar reference cell type.

We averaged and compared pairwise expression distances (Figure S7) for different groups (Figure 4D): the distances among iPSC clones within and between each species (N=14 samples), the average of the distances between 1383D2 and the urinary derived human iPSCs (N=9) and the average of the pairwise distance between and within individuals among iPSCs and species (within individuals: N=6 (6 individuals with more than one clone), between individuals: N=8).

## Acknowledgments

We thank Stefanie Färberböck for her expert technical assistance and enormous help in cell culture. We are grateful to Christine Gohl and the staff at the Zoo Hellabrunn for kindly collecting and providing the primate urine samples.

## Author Contributions

J.G., M.O. and W.E. conceived the study. J.G. and W.E. wrote the manuscript. J.G. established iPSC lines and conducted differentiation experiments. J.G., L.E.W. A.J. and J.W.B. prepared sequencing libraries and processed data. A.K. tested virus absence in primary cells. S.M. and J.G. performed karyotype analyses of iPSC lines.

