## Supplemental Figures for "A non-invasive method to generate induced pluripotent stem cells from primate urine"

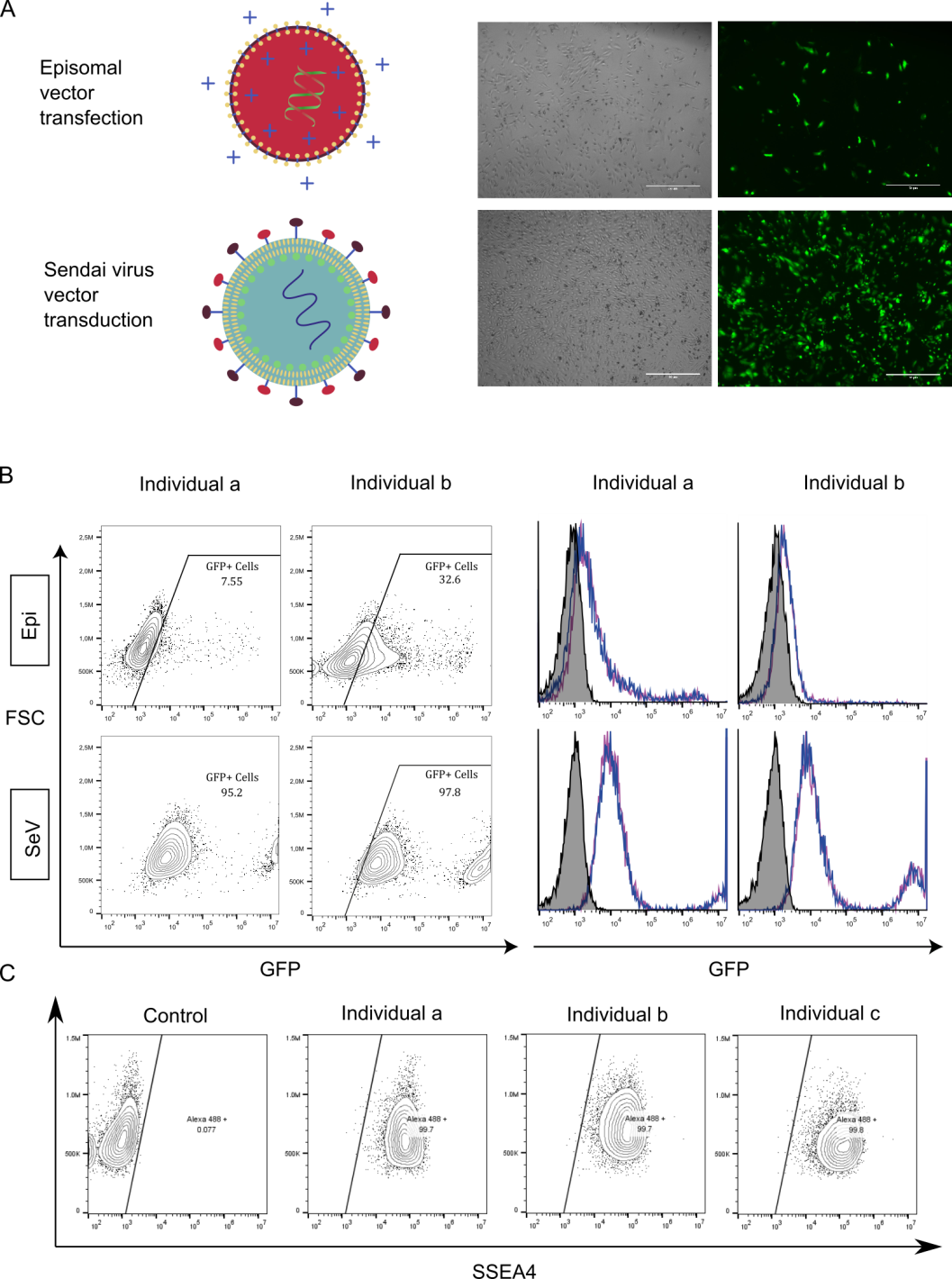

**Figure S1. Transfection/Transduction efficiency of urinary cells, Related to Figure 1**

**(A)** GFP expression of urinary cells transfected with pcxle-EGFP episomal plasmids or CytoTune EmGFP transduced after 5 days **(B)** FACS analysis of GFP expressing cells 5 days post transfection/transduction **(C)** SSEA4 expression of urinary cells

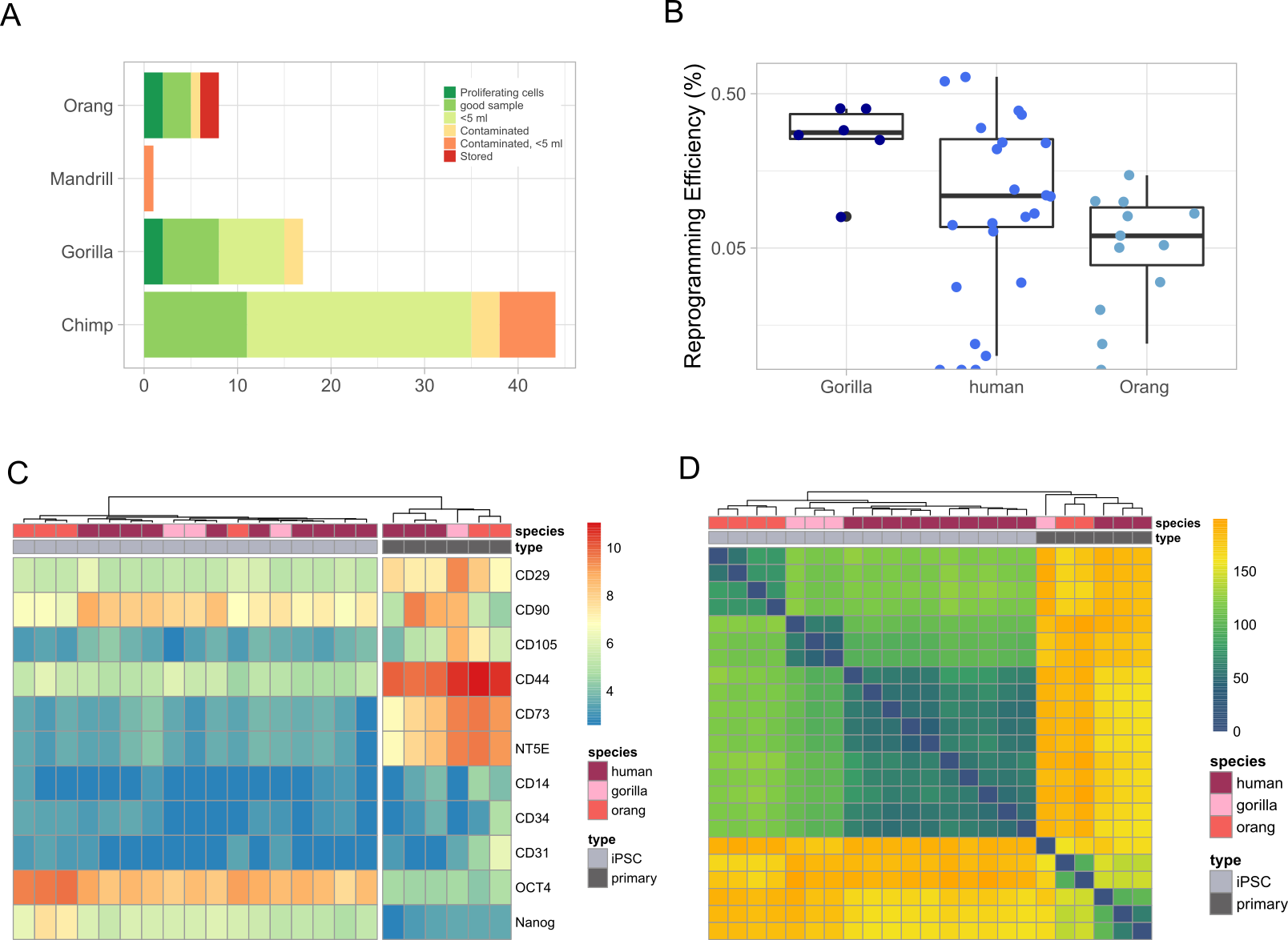

**Figure S2. Urine isolation and reprogramming efficiencies, Related to Figure 2 and Figure 4**

**(A)** Overview of collected urine samples and properties of the samples, associated to successful isolation of proliferating cells. **(B)** Reprogramming efficiency shown as colonies per number of seeded cells between species. **(C)** Heatmap of mesenchymal stem cell and iPSC marker expression. **(D)** Euclidean distance between samples.

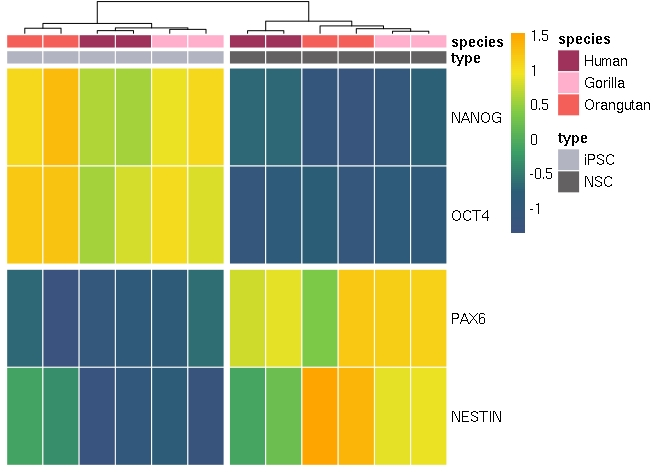

**Figure S3. Marker expression in iPSCs and NPCs, Related to Figure 3**

Dual-SMAD inhibition leads to the formation of neurospheres in floating culture, confirmed by neural stem cell marker expression (NESTIN+, PAX6+) using qRT-PCR.

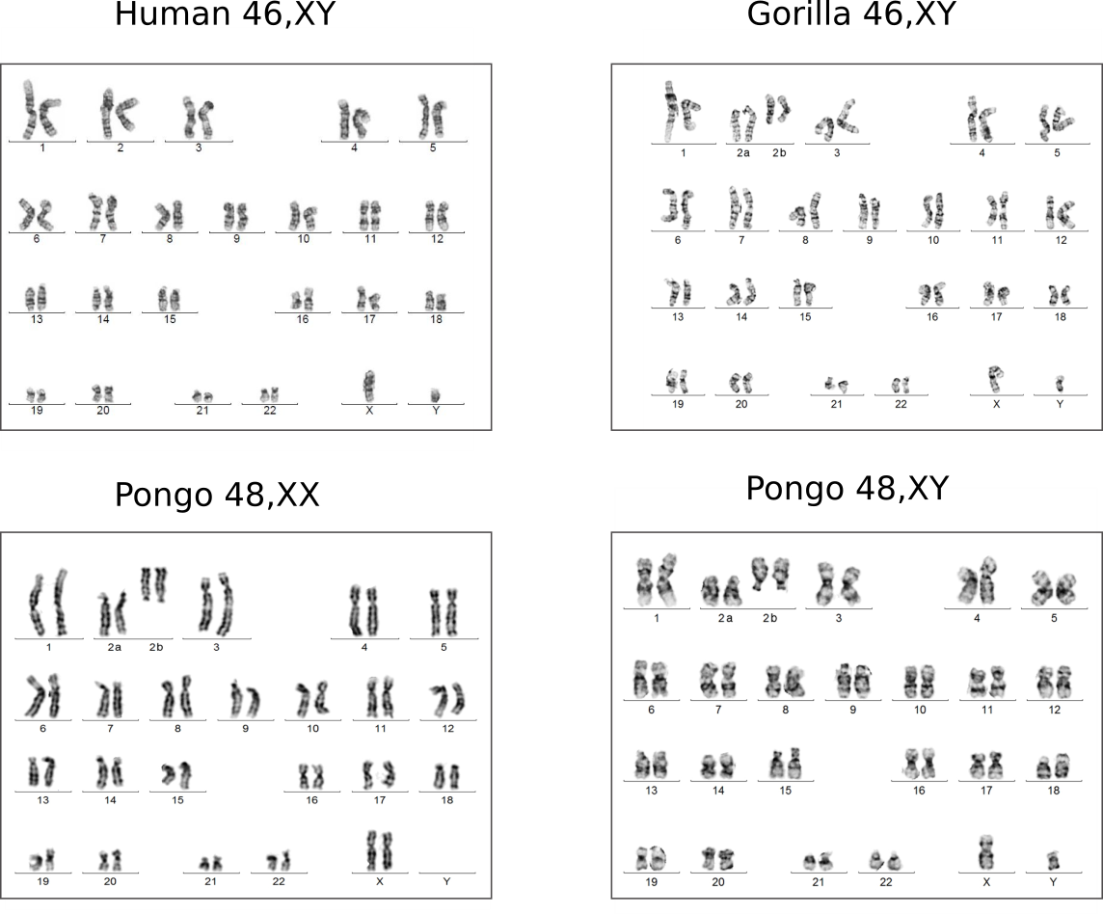

**Figure S4. Karyograms of primate iPSC lines, Related to Figure 3**

Exemplary karyotyping analysis of Primate iPSCs. Five out of five tested iPSC lines show a normal karyotype without chromosomal alterations.

**Table S4. Primers used for qPCR, Related to Experimental Procedures**

|  | Forward | Reverse |
| --- | --- | --- |
| SeV | GGA TCA CTA GGT GAT ATC GAG C | ACC AGA CAA GAG TTT AAG AGA T |
| GAPDH | ACC ACA GTC CAT GCC ATC AC | TCC ACC ACC CTG TTG CTG TA |
| hOCT3/4 | GAC AGG GGG AGG GGA GGA GCT AGG | CTT CCC TCC AAC CAG TTG CCC CAA AC |
| NESTIN | GCC CTG ACC ACT CCA GTT TA | GTC CTG GAT TTC CTT CC |
| PAX6 | CTT GGG AAA TCC GAG AGA GA | CTA GCC AGG TTG CGA AGA AC |
| NANOG | GAT TTG TGG GCC TGA AGA AA | CAG ATC CAT GGA GGA AGG AA |
